# Comethyl: A network-based methylome approach to investigate the multivariate nature of health and disease

**DOI:** 10.1101/2021.07.14.452385

**Authors:** Charles E. Mordaunt, Julia S. Mouat, Rebecca J. Schmidt, Janine M. LaSalle

**Affiliations:** Department of Medical Microbiology and Immunology, Genome Center, Perinatal Origins of Disparities Center, and MIND Institute, University of California, Davis, CA, USA; Department of Public Health Sciences, Perinatal Origins of Disparities Center, and MIND Institute, University of California, Davis, CA, USA

**Keywords:** DNA methylation, whole genome bisulfite sequencing, epigenetics, epigenome, weighted gene correlation network analysis, systems biology, autism spectrum disorder

## Abstract

Health outcomes are frequently shaped by difficult to dissect inter-relationships between biological, behavioral, social, and environmental factors. DNA methylation patterns reflect such multi-variate intersections, providing a rich source of novel biomarkers and insight into disease etiologies. Recent advances in whole-genome bisulfite sequencing (WGBS) enable investigation of DNA methylation over all genomic CpGs, but existing bioinformatic approaches lack accessible system-level tools. Here, we develop the R package Comethyl, for weighted gene correlation network analysis (WGCNA) of user-defined genomic regions that generates modules of comethylated regions, which are then tested for correlations with sample traits. First, regions are defined by CpG genomic location or regulatory annotation and filtered based on CpG count, sequencing depth, and variability. Next, correlation networks are used to find modules of interconnected nodes using methylation values within the selected regions. Each module containing multiple comethylated regions is reduced in complexity to a single eigennode value, which is then tested for correlations with experimental metadata. Comethyl has the ability to cover the noncoding regulatory regions of the genome with high relevance to interpretation of genome-wide association studies and integration with other types of epigenomic data. We demonstrate the utility of Comethyl on a dataset of male cord blood samples from newborns later diagnosed with autism spectrum disorder (ASD) versus typical development. Comethyl successfully identified an ASD-associated module containing gene regions with brain glial functions. Comethyl is expected to be useful in uncovering the multi-variate nature of health disparities for a variety of common disorders. Comethyl is available at github.com/cemordaunt/comethyl.

**Description of the Authors:** **Charles E. Mordaunt**, Ph.D. developed Comethyl while a postdoctoral fellow in the department of Medical Microbiology and Immunology at UC Davis. He is currently a Computational Biologist at GSK.

**Julia S. Mouat** is a doctoral student in the Integrative Genetics and Genomics graduate group at UC Davis with interests in health disparities and intergenerational epigenetic risk factors for autism spectrum disorders.

**Rebecca J. Schmidt**, Ph.D. is an Associate Professor of Public Health Sciences at UC Davis, with expertise in the use of epigenetics in epidemiology and neurodevelopmental disorders.

**Janine M. LaSalle**, Ph.D. is a Professor of Medical Microbiology and Immunology, Co-Director of the Perinatal Origins of Disparities Center, and Deputy Director of the Environmental Health Sciences Center at UC Davis, with expertise in epigenomics and neurodevelopmental disorders.

## Introduction

Despite the exceptional promise of genetics in understanding human health and disease, individual genes do not act in isolation from each other or from outside influences [1]. For instance, human susceptibility to SARS-COV2 is highly variable due to behavioral, social, and economic factors influencing exposures in addition to underlying health disparities and genetics [2][3]. Recent advances in genome sequencing technologies have greatly expanded our potential to understand the complex relationships that influence biobehavioral health trajectories and outcomes. However, existing statistical and analytical tools are frequently overly reductionist and specialized at determining the role of a single gene or environmental factor in disease by making precarious assumptions about the lack of gene x environmental interactions [4]. Unlike the relatively static genome or the highly dynamic transcriptome, the DNA methylome is poised at the interface of genetics and a variety of environmental influences, reflecting past cellular variations in gene expression [5]–[8]. Through whole-genome bisulfite sequencing (WGBS), the DNA methylome is becoming a tractable “gold mine” for discovering novel biomarkers and improved insights into disease mechanisms [9]. What is currently lacking for WGBS and other DNA methylation technologies, however, is an improved analytical pipeline that intersects gene-level investigations with systems-level integration of multivariate data.

We developed the user-friendly R package Comethyl, available at github.com/cemordaunt/comethyl, for integration of WGBS data with additional metadata related to experimental conditions, cell type heterogeneity, and patient demographics. The weighted gene correlation network analysis (WGCNA) R package [10] was designed to perform network analysis of transcriptome data, so the gene is the unit of analysis, but transcribed genes make up only ~2% of the human genome. Additionally, WGBS data requires appropriate aggregation and filtering methods that differ from gene expression data. In Comethyl, we therefore developed an interface that extends WGCNA to allow user-determined genomic regions of clustered CpGs to be investigated for interconnectedness. Regional clusters of CpGs are frequently correlated in methylation levels and are biologically more informative than individual CpGs because they are less susceptible to stochastic variation. Regions can also be defined based on other functional annotations, including gene bodies, promoters, and enhancers. Once genomic regions are selected, the Comethyl package then adapts the existing WGCNA approach to identify modules of interconnected genomic regions based on correlated methylation levels. Once assigned to modules, these genomic regions can be mapped to genes, explored for enrichments to regulatory annotations and gene functions, and correlated with individual traits from experimental metadata to provide insights into complex conditions.

Autism spectrum disorder (ASD) is a complex, heterogeneous genetic condition with predicted gene x environmental interactions in perinatal life [11][12]. Using a high-risk prospective human cohort, we recently published a cord blood WGBS analyzed using a differentially methylated region (DMR) approach that identified novel ASD associated loci [13]. In order to test the utility of Comethyl in identifying disease-relevant associations, we re-analyzed male cord blood WGBS data and identified a novel module that was negatively correlated with later ASD diagnosis but not with other cell type, demographic, or experimental factors. Here, we outline our analytical approach in Comethyl applied to this real-world dataset and discuss how it can be applied more broadly.

## Methods and Results

### Comethyl pipeline, data requirements, and CpG filtering

The Comethyl pipeline has three major subsections, shown as columns in the flow chart in Figure 1. The first section involves loading cytosine reports as well as defining CpGs and regions to be included in downstream analyses. Principal components of the included regions are identified and automatically adjusted for. Next, a comethylation network is constructed to call comethylation modules and characterize sample and module correlations. Lastly, modules are tested for significant correlations with relevant traits and experimental variables, mapped to genes, and assessed for functional enrichments.

**Figure 1.**
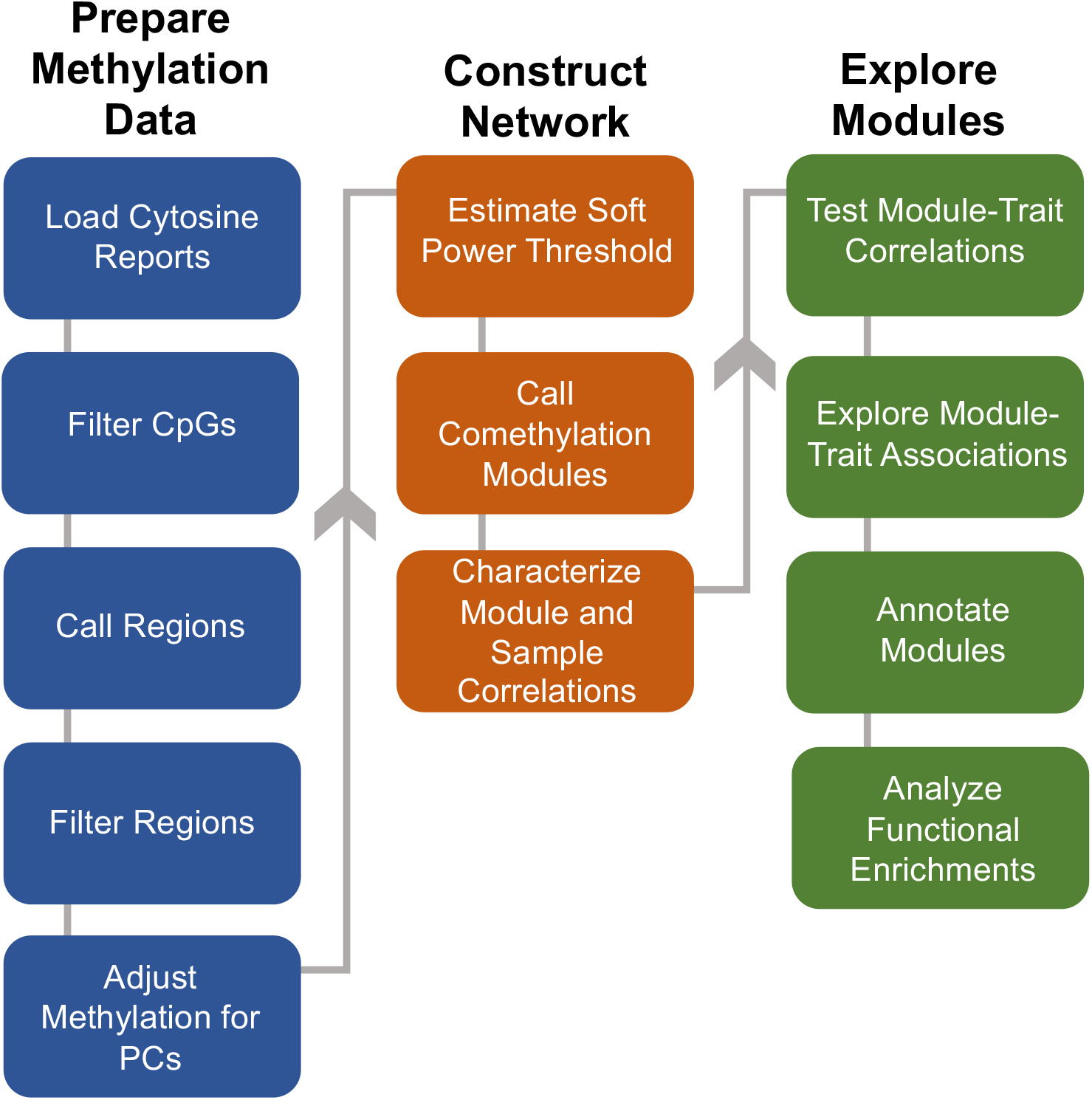
Comethyl pipeline for weighted region comethylation network analysis. Starting with CpG-level data from whole-genome bisulfite sequencing, the Comethyl pipeline consists of a suite of functions that allow users to summarize DNA methylation at the region level in a biologically-driven manner, construct a comethylation network, and explore associations of comethylation modules with traits.

We explored the use of Comethyl in a previously published WGBS dataset of 74 male newborn cord blood samples, including those that were diagnosed with ASD or classified as typically developing by 36 months of age (TD *n* = 39, ASD *n* = 35). WGBS read alignment and quality control was performed using the CpG_Me workflow [14]–[17], which includes the generation of a Bismark CpG report. In Comethyl, individual sample Bismark CpG reports are read into a single BSseq object. The user can then select cutoff values to filter the CpGs. The cutoffs pertain to sequencing coverage and percent of samples, both of which should be balanced with the total number of CpGs included in downstream analysis (Figure S1A, Table S1). With low-pass WGBS (3-8x average CpG coverage), using strict or invariant coverage thresholds greatly reduces the number of genomic CpGs assayed, particularly as sample size increases. In this dataset, CpGs were filtered to include those with a minimum 2x coverage in 75% of the samples in order to balance total CpG number with sufficient sequencing depth and sample representation. Because Comethyl focuses on region-level methylation, relatively sparse CpG-level counts are aggregated together into regions for improved accuracy, and high coverage at individual CpGs is less critical.

### Calling and filtering of genomic regions

Since DNA methylation of individual CpG nucleotides is imprecise but clusters of CpGs are regionally correlated, Comethyl summarizes CpG methylation at the region level prior to WGCNA. In the ASD cord blood data set, regions are defined as CpG clusters containing at least 3 CpGs separated by no more than 150 bp, though this can be adjusted by the user. Region features, such as the total number of regions, the total width of regions, and total CpGs, are then visualized in comparison with minimum coverage and methylation standard deviation cutoffs to aid the user in determining appropriate thresholds for region filtering (Figure S1B, Figure 2, Table S2). In the case of the ASD cord blood data, regions were selected for those with at least 10 reads in all samples and a standard deviation greater than 0.05, resulting in 251,717 regions for further analysis (Table S3). The overall goal of this step is to reduce the total number of regions assayed to those with sufficient coverage and variation in methylation between samples, which are thus the most likely to exhibit interconnectivity and informative relationships with traits of interest.

**Figure 2.**
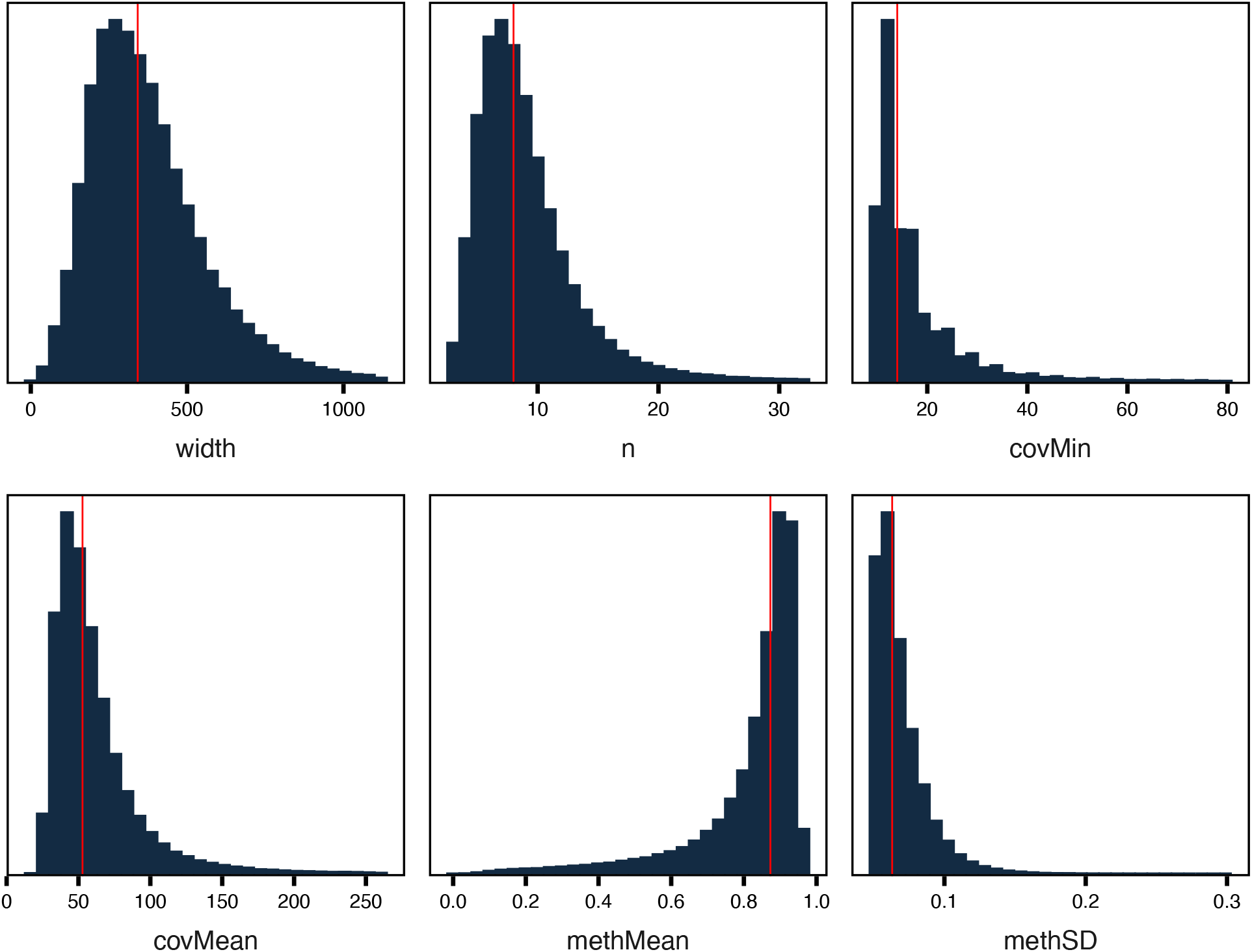
After filtering of CpG clusters, most regions have similar characteristics. CpGs with sufficient coverage in WGBS were grouped into clusters of at least 3 CpGs separated by no more than 150 bp and filtered for those with at least 10 reads in all samples and standard deviation greater than 0.05. Plots show the distributions of multiple region characteristics with the red line indicating the median value.

In addition to the approach of calling genomic regions by CpG location, Comethyl also includes the option to define regions based on functional annotations, such as CpG islands, gene bodies, enhancers, or a custom annotation. Since gene body methylation can correlate positively with expression [18], [19], gene bodies were selected as alternative regions to explore in the same cord blood dataset. Similar to the analysis of regions, gene bodies were then filtered for those with at least 3 CpGs and 10 reads in all samples (Figure S2, Table S4, Table S5).

### Preparation for network construction

Before building the comethylation network, it is necessary to adjust the methylation data for confounding factors, check for outlier samples, and evaluate scale-free topology. By default, percent methylation is directly calculated as the total methylated reads divided by the total reads in a region. One option is to instead use smoothed methylation data, for example using the BSmooth algorithm, which may be advantageous for low coverage datasets. Once the methylation values are obtained, Comethyl uses an unbiased principal component approach to adjust for the dominant variables in the data. This approach adapts sva_network() from the sva R package and is specifically designed to remove confounding technical variables and still allow for downstream network analysis. The required number of principal components is estimated and then the methylation data is adjusted. In the case of the cord blood data, the top 10 principal components were used for adjustment.

In order to check for outlier samples, the adjusted methylation data is used to cluster samples by Euclidean distance (Figure S3A for regions, S4A for gene bodies). Next, the scale-free topology is assessed with the goal of selecting a soft power threshold for network construction. The soft power is the power to which all correlations between genes are raised, a process that decreases background noise from weak correlations and amplifies stronger correlations. Soft power is plotted against the scale-free topology fit and mean connectivity, which is the number of edges between nodes in the network and is inversely correlated with fit. It is recommended to select a soft power threshold that provides a fit of at least 0.8. The scale-free topology can be assessed with Pearson or biweight midcorrelation (Bicor) statistics to correlate genes. Pearson correlation is mean-based and more sensitive to outliers because it assumes that methylation data follows a normal distribution. In contrast, Bicor correlation is median-based and thus less sensitive to outliers; however, it also has reduced power to detect correlations. Because of the increased power with Pearson correlation, we used this for soft power estimation and network construction in the cord blood dataset. For datasets with high variability in methylation values, it may be more appropriate to use Bicor instead. It can be helpful to examine the modules and trait correlations observed for different network methods, in order to select the method that best fits the particular dataset. For the analysis of regions in the cord blood dataset, a soft-thresholding power cutoff of 18 was selected as the lowest power with a fit of at least 0.8 (Figure S3B). For the analysis of gene body-defined regions, a soft power threshold of 12 was selected (Figure S4B).

### Network construction and calling of Comethylation modules

The next step in Comethyl is to identify distinct groups of regions with correlated methylation values, defined as “comethylation modules”. Using the chosen soft power threshold and statistical model from the previous step, module construction uses a two-phase clustering approach to reduce computational intensity, which is implemented using the WGCNA package. In the first phase, regions are formed into large blocks using k-means clustering. In the second phase, a full network analysis is performed on each block to assess the module membership of each region. Similar modules from different blocks can then be merged. Figure 3A shows the dendrogram with color-coded modules identified from one of eight blocks formed from the cord blood dataset. Regions not assigned to comethylation modules are assigned to the grey module. Interestingly, regions assigned to a module make up a minority of cases. Regions within each module are distributed throughout the genome and may overlap multiple genes and/or CpG islands (Figure 3B). In this dataset, 53 modules were detected, and these ranged from the thistle2 module with 10 regions to the turquoise module with 61 regions (Figure 3C, Table S3). With the gene body-based regions, 15 modules were identified, each including 10-139 regions (Figure S5, Table S5).

**Figure 3.**
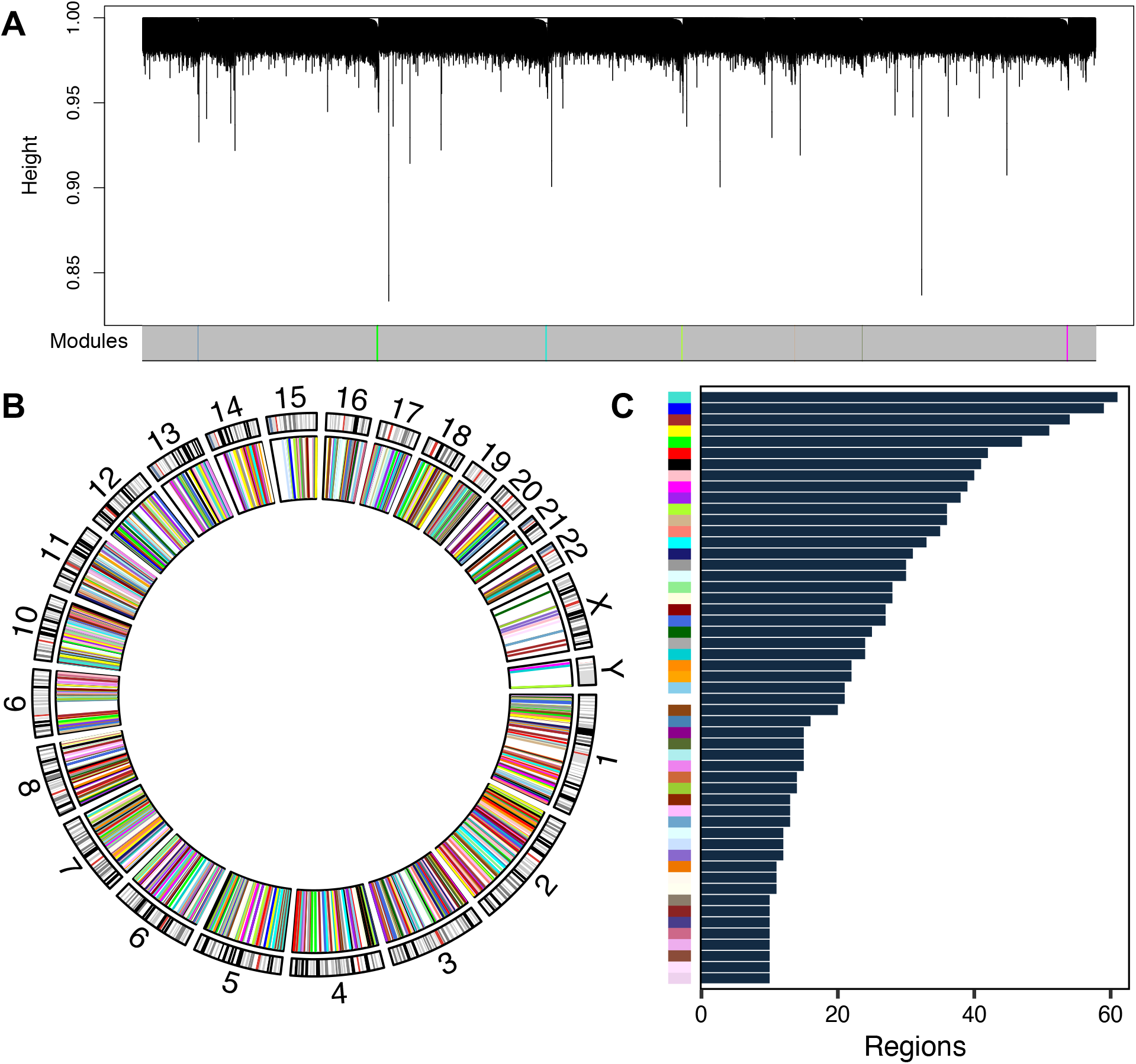
CpG cluster comethylation modules identified in cord blood. A comethylation network was constructed from adjusted region methylation data and assessed for modules of comethylated regions. (A) Region dendrograms for one block. (B) Circos plot of regions colored by module in the inner ring and chromosome bands in the outer ring. (C) Number of regions assigned to each module.

Overall, we expect only a small number of regions (<1-2%) to be assigned to a module, at least for samples from a single tissue or cell type that come from a population with a complex disease. In contrast, a large portion of regions being assigned to a single module could indicate a confounding variable that has not been addressed with principal component adjustments. Correlations of modules with samples and traits may reveal additional information to guide improved normalization.

### Characterization of module and sample correlations

Eigennode values, which summarize the methylation values of each module, are calculated for each sample and then used to further examine relationships between samples and modules (Figure S6, S7). An expected dynamic range of eigennode values was observed in the cord blood data, with samples clustering on the basis of higher or lower eigennode values of neighboring modules. Heatmaps of between-sample and between-module (Figure 4, S8) correlations of eigennode values are also generated, revealing patterns of interconnectedness in the methylation data. Similarities based on module methylation between samples and modules can be investigated, potentially revealing subsets of related samples or modules.

**Figure 4.**
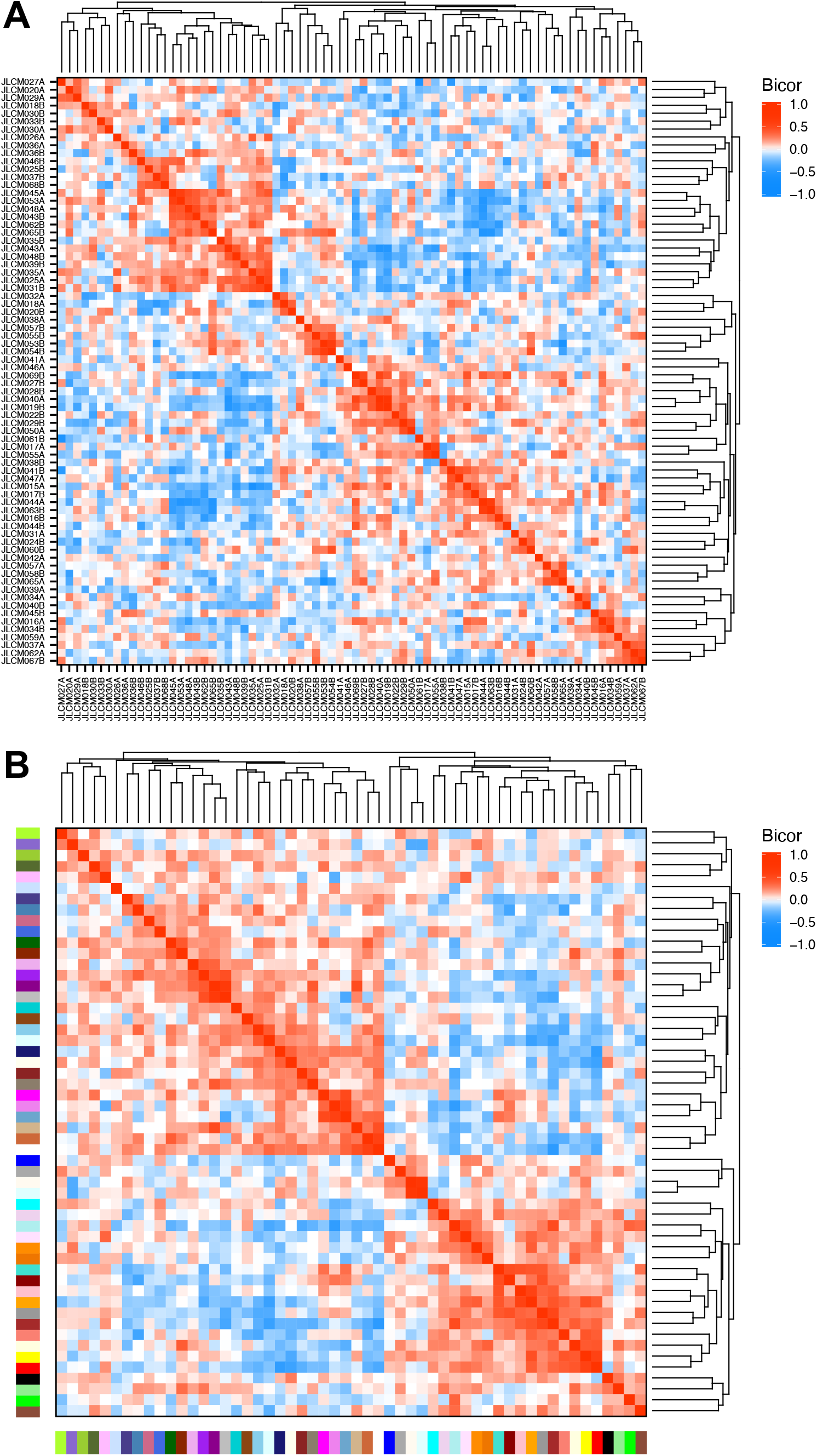
Correlations of CpG cluster module eigennode values reveal similarities between subsets of samples and modules. (A) Eigennode values were clustered and compared across samples using biweight midcorrelation. (B) Same as (A), but compared across modules.

### Test all module-trait associations and explore informative modules

Once comethylation modules are identified and eigennode values defined for each module and sample, both continuous and discrete traits of the samples can be explored for module-trait associations using Pearson or Bicor correlations. Sample traits can include all available information about potential variables of interest as well as potential confounding variables. In WGBS datasets, potential confounding variables include cell type proportions as well as technical variables including coverage, read duplication, read trimming, and global cytosine methylation levels. For human populations, metadata should include clinical, diagnostic, and demographic data, as well as sample collection characteristics, such as gestational age and birthweight for cord blood. For experimental studies in animal models or cell cultures, experimental variables should be included in the metadata for exploring module-trait relationships. One of the main features of this system-level approach is that correlations with many traits can be assessed simultaneously and then explored in greater detail.

Module-trait Bicor correlations between all 49 variables and 53 modules from the unbiased region analysis in the cord blood dataset were tested (Figure 5A, Table S6). After false discovery rate *p*-value adjustment, a small number of module-trait associations were significant (FDR *p* < 0.05, Figure 5B). One comethylation module (Bisque 4) showed a significant negative correlation between sample eigennode values and ASD diagnosis, but none of the other potential associations with this module were significant after correction. Cord blood samples from newborns later diagnosed with ASD showed lower Bisque 4 module eigennode values than those who were diagnosed as typically developing (Figure 5C). Several other comethylation modules were significant for associations with one or more traits, most frequently cell type proportions (granulocyte, B cell, nucleated red blood cell, CD4+ T cell) but also some technical variables (whole-genome % CpG or CpH methylation, or cohort study site). Assessing a range of variables is useful in determining which modules are of the greatest interest for further investigation, and which are confounded by other factors.

**Figure 5.**
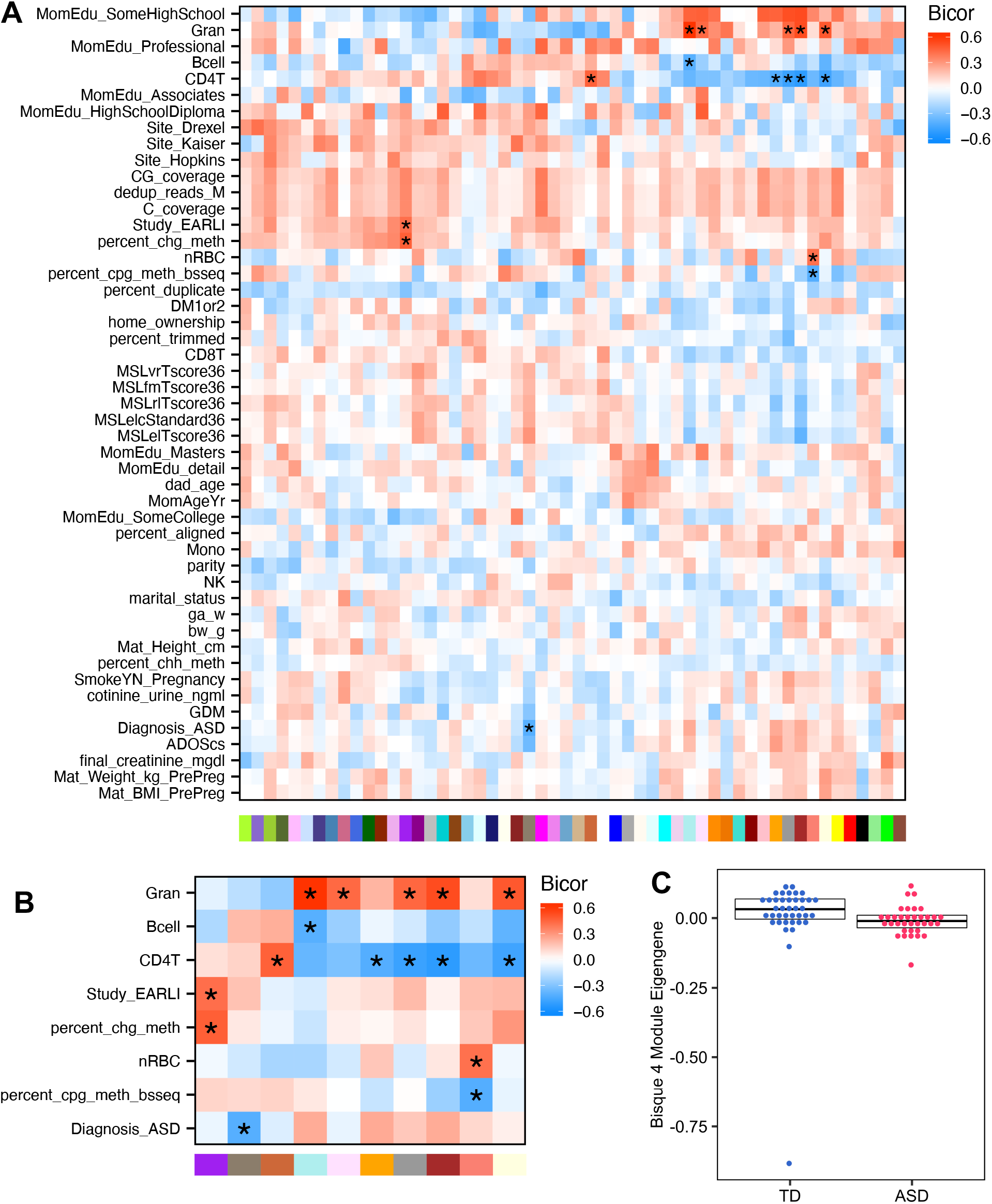
Correlations of CpG cluster module eigennodes with sample traits. (A) All tested correlations between module eigennode values and sample traits using biweight midcorrelation. (B) Same as (A) but significant modules and traits only (* FDR-adjusted *p* < 0.05). (C) Bisque 4 module is associated with later ASD diagnosis. Eigennode values are plotted for each sample in relation to ASD diagnosis. Box indicates 1^st^ quartile, median, and 3^rd^ quartile.

In the heatmap generated from the gene body-defined regions of the same cord blood dataset, no modules were significant for association with ASD diagnosis after FDR correction, although several modules were significant for cell type proportions or whole-genome CpG methylation (Figure S9, Table S7). The Green-Yellow module was significantly associated with the discrete demographic variable of Home Ownership, as well as the continuous variable of cell type percentage for granulocytes, CD4+ T cells, and CD8+ T cells (Figure S10). More specifically, higher Green-Yellow module eigennode values were associated with home ownership, lower levels of granulocytes, and higher levels of CD4+ and CD8+ T cells. Notably, association with cell type proportion does not necessarily exclude a module from further consideration, but it does add important biological context for the interpretation of results.

### Annotate modules and test for functional enrichments

The last step in Comethyl is to annotate genes to module regions and test enrichment of genes within each module for specific functions. For modules defined by the analysis of unbiased genomic regions, genes are annotated using Genomic Regions Enrichment of Annotations Tool (GREAT), gene information is added from BioMart, and gene context and CpG island context is added from annotatr. The Bisque 4 module contains 10 regions, which map to 17 genes because most are in intergenic regions and all are in “open sea” locations with respect to CpG islands (Table 1). For gene body-defined regions, genes are by definition already annotated. Table 2 shows an example of the output table from the Green-Yellow module associated with home ownership, containing 14 gene bodies.

**Table 1.** Regions and nearby genes identified in the Bisque-4 module associated with ASD diagnosis.

| chr | start | end | CpGs | hubRegion | gene_symbol | distance_to_TSS | gene_context | CpG_context |
| --- | --- | --- | --- | --- | --- | --- | --- | --- |
| chr1 | 57479875 | 57479935 | 7 | TRUE | *DAB1* | -55848 | intron | open sea |
| chr2 | 224079272 | 224079787 | 9 | FALSE | *SERPINE2,*<br>*FAM124B* | -40726, 322504 | intergenic | open sea |
| chr5 | 94040899 | 94041136 | 7 | FALSE | *POU5F2,*<br>*FAM172A* | -299381, 70617 | intron | open sea |
| chr6 | 47424495 | 47424763 | 4 | FALSE | *TNFRSF21,*<br>*CD2AP* | -114724, -53160 | intergenic | open sea |
| chr10 | 35134739 | 35134967 | 7 | FALSE | *CCNY, CREM* | -202021, 7393 | promoter,<br>1to5kb, intron | open sea |
| chr11 | 80720879 | 80721186 | 14 | FALSE | NA | NA | intergenic | open sea |
| chr13 | 52925862 | 52926068 | 11 | FALSE | *OLFM4,*<br>*PCDH8* | -102794, -77325 | intergenic | open sea |
| chr15 | 68240587 | 68240932 | 12 | FALSE | *FEM1B, CLN6* | -37043, -11017 | intron | open sea |
| chr15 | 99715416 | 99715665 | 8 | FALSE | *LYSMD4,*<br>*MEF2A* | 17880, 149454 | exon, intron,<br>3UTR | open sea |
| chr19 | 43838028 | 43838499 | 6 | FALSE | *ZNF283,*<br>*ZNF404* | 10972, 45700 | exon, intron,<br>3UTR | open sea |

**Table 2.**
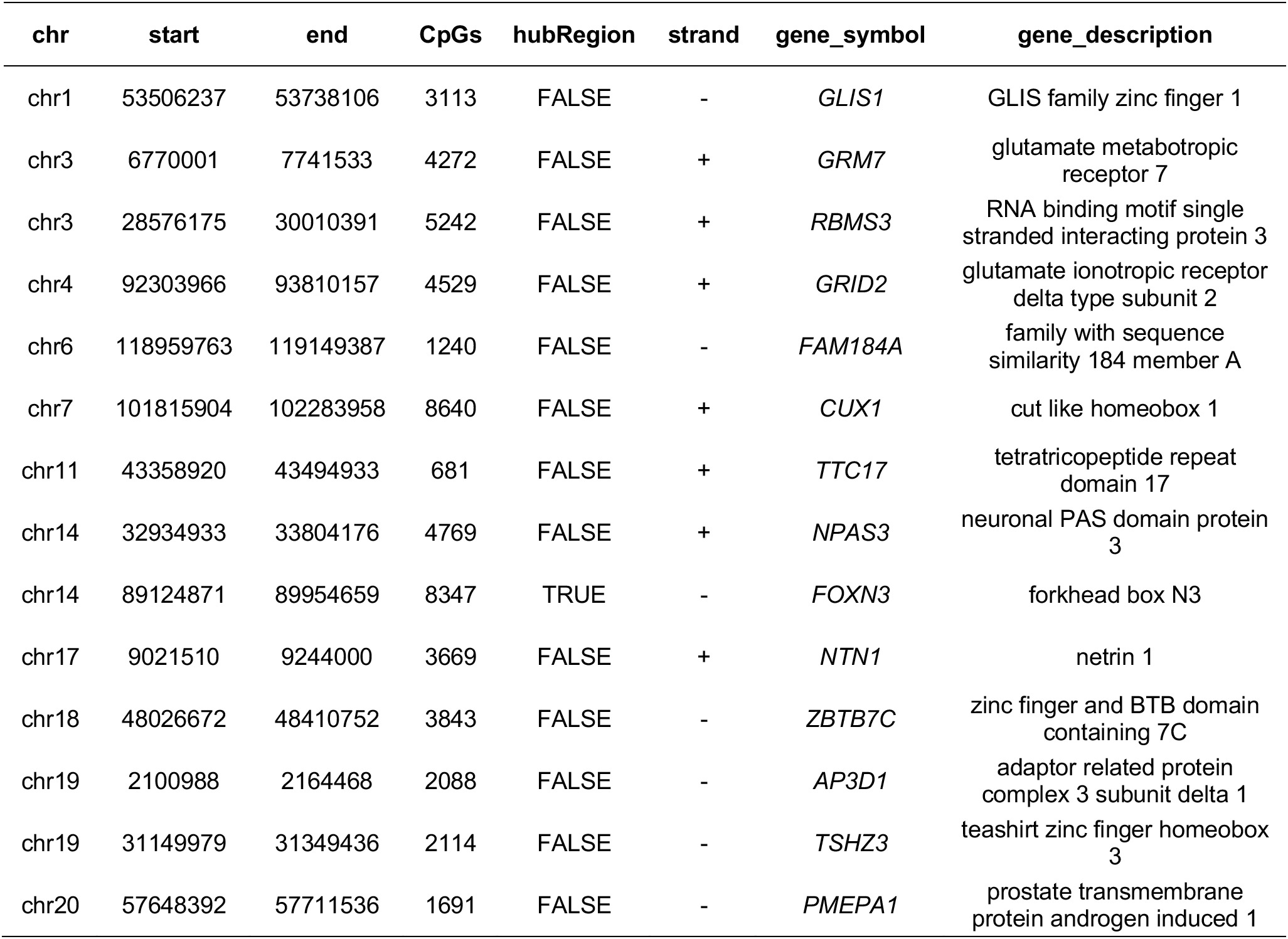
Gene bodies within the Green-Yellow Module associated with home ownership and three cell types.

Functional enrichment analysis in GREAT is performed on regions within each module compared to all regions tested and examined for overrepresentation of genes in Gene Ontology biological functions, cellular components, and molecular functions, along with both human and mouse phenotype-associated genes. For the Bisque 4 module associated with ASD diagnosis, genes were enriched in functions related to glial cells and glycosphingolipid metabolism, relevant to cerebellar development and thus ASD (Figure S11, Table S8). For the Green-Yellow gene body module associated with home ownership and immune cell types, the 14 genes were enriched in glutamate receptor activity and several abnormal phenotypes, including abnormal respiration, postnatal lethality, and increased prepulse inhibition (Figure S12, Table S9). Functional enrichment can be used to characterize the potential regulatory role of comethylation modules.

## Discussion

Comethyl was designed as a user-friendly R package for analyzing WGBS data from human studies or relatively complex experimental animal models where multiple variables are potentially associated with or interconnected in influencing DNA methylation levels. While WGCNA approaches have been utilized in the analysis of genome-wide DNA methylation previously [20][21][22][23][24][25][26][27], Comethyl appears to be the first one that utilizes the sequencing-based merits of WGBS data. In order to provide the most unbiased coverage of all genomic regions, including those over poorly annotated intergenic regions, Comethyl is specifically designed to empirically define and filter the regions that go into downstream module assignments. In addition to identifying regions agnostic to functional annotations, the Comethyl pipeline includes the flexibility for a user to conduct a focused analysis through the selection of gene bodies, CpG islands, or other annotations as regions. A user could additionally integrate any type of region-based annotation with Comethyl, such as those from ChromHMM chromatin segmentations or ATAC-seq open chromatin peaks.

Comethyl was designed to work predominantly on low-pass WGBS data based on regional groups of CpGs similar to DMRs. If Comethyl is used to analyze EPIC/450k data, it is recommended to create regions of clustered CpGs or limit analysis to gene promoters or gene bodies. Alternatively, an approach developed for array data, such as CoMeBack, could be used [28]. The major limitation with array data is the sparse representation of CpGs over most of the genome and the difficulty in creating informative regions from biased CpG sampling [29], [30]. For instance, several gene modules were associated with ASD diagnosis in a human brain study, but the gene functions identified were primarily associated with immune function [20]. In contrast, our sequencing-based analysis of cord blood samples from newborns who were later diagnosed with ASD identified brain development and glial functions. Additionally, the region-level approach in Comethyl aggregates low-pass WGBS CpG-level data into regions to improve accuracy without requiring excessive sequencing depth. Since the majority of WGBS-covered regions lie outside of those covered on EPIC and 450k arrays, they have been missed by array-based methylome studies. To avoid the inevitable health disparities that can arise from the exclusive use of genomic platforms designed to human subjects of European ancestry [31], [32], sequencing-based approaches including WGBS and Comethyl offer the ability to make discoveries with broader relevance to human populations of diverse ancestry as well as to genomic regions that are poorly annotated.

Comethyl was designed to be complementary to DMR-based approaches to understand DNA methylation differences associated with diseases and exposures. Comethyl may be used in conjunction with DMR analysis to examine the correlated nature of CpG sites and their functional enrichments. The strength of DMR analyses is that they are more likely to identify specific individual genetic loci of high significance. However, the weakness is that some potential biologically relevant DMRs could be lost to noise in the underlying data and interacting variables affecting methylation levels. In contrast to DMR analysis, Comethyl does not consider absolute methylation values of the samples to correlate CpG sites/regions. Instead, Comethyl allows the user to set windows or group adjacent CpGs, within which directional changes in methylation are used to group regions into modules. By correlating user-defined windows of the genome, Comethyl can analyze samples that have different absolute methylation values, which is prevented by other approaches such as DMR analysis. Through identifying sets of regions whose methylation varies together across samples, Comethyl may also identify regions with functionally related genes and link them to a sample trait. Another major advantage of Comethyl over DMR approaches is the ability to simultaneously examine all potential confounding associations with technical, experimental, and demographic variables with the same comethylation modules as those with traits of interest. It is therefore recommended that Comethyl precede DMR analyses of WGBS data in complex datasets, so that confounding variables can be identified and adjusted for in DMR calling. Traits can then be prioritized for those most likely to be associated with altered DNA methylation. Further, Comethyl has the potential to reveal important insights into health disparities in the data, such as the association between home ownership, considered a surrogate of socioeconomic status, and T cell and granulocyte proportions with the methylation at a set of genomic regions, which was observed in our analysis.

Integration of DNA methylation data with other large-scale multi-omics datasets is a major challenge for the future [33]. Comethyl reduces the complexity of the methylation analysis of 29 million CpG sites assayed to 20-80 distinct modules for which a single eigennode value is assigned per sample. In this way, comethylation modules associated with the disease or trait of interest can be compared to additional measurements of phenotypes, metabolites, and microbiota. Comethyl can be used in conjunction with gene expression analysis by comparing the gene body methylome with the transcriptome in the same sample population. This comparison allows for greater insight into epigenetic regulation of gene pathways and their role in disease, as well as the validity of epigenetic biomarkers in various biological contexts. Because Comethyl covers both coding regions and noncoding regulatory regions of the genome, it is highly relevant to the interpretation of common human polymorphisms identified from genome-wide association studies (GWAS). Genes and genomic regions annotated to comethylation modules that are significantly correlated with a trait of interest, such as diagnosis, can be compared to those identified by GWAS for potential overlap and examined for genetic effects on DNA methylation. Lastly, regions defined for Comethyl analysis can be defined from additional layers of epigenetic information, such as chromatin state maps (chromHMM) defined from histone modifications, open chromatin, and chromatin loop binding sites.

There are several potential limitations of Comethyl that should be considered during analysis. First, since most regions map distal to known genes, there is only an indirect link to the closest mapped genes, which may not be those functionally influenced by the methylation changes. Additional chromatin contact datasets or follow up experiments are needed to confirm gene mappings. Second, some biologically relevant methylation changes may be missed in regions filtered out based on low CpG density, low methylation variability, or overly aggressive principal component adjustments. Therefore, a complementary approach of both Comethyl and DMR analyses is recommended to provide validation of the most reproducible effects, as well as to balance system-level with targeted analysis and find important associations that were missed with the alternate approach.

Future goals for Comethyl include adding measured environmental exposures, polygenic risk scores, and circadian rhythmicity as traits to be correlated with modules. Including specific environmental exposures as traits would allow for direct correlation with epigenetic patterns regulating gene modules, as well as elucidating potential gene by environment interactions. Examining the functional enrichments of modules significantly correlated with exposures of interest could further suggest potential mechanisms of action. Adding polygenic risk score as a trait in a study of complex disease would allow for integrated analysis of genetic risk, environmental variables, and epigenetic patterns. It would be enlightening, for example, if polygenic risk and a specific environmental variable were both significantly associated with a comethylation module. This could generate hypotheses about gene-environment interactions and disease etiology that could be explored in later functional studies.

### Key Points

- We developed an R package, Comethyl, to bring multiple tools together and enable comethylation network analysis from low-pass WGBS data, which is available at github.com/cemordaunt/comethyl.
- Comethyl is a complete workflow, from CpG-level counts to network construction, to functional enrichment of comethylation modules.
- We applied Comethyl to an ASD cord blood dataset with 2 different region methods and identified comethylation modules associated with traits of interest, including ASD, and enriched in relevant biological functions.
- Comethyl allows for system-level investigation into the impact of diverse traits on the methylome and on gene regulation.

## Data Availability

The data underlying this article are available in the Gene Expression Omnibus at https://www.ncbi.nlm.nih.gov/geo/ and can be accessed with accession number GSE140730.

## Funding

This work was supported by the National Institutes of Health R01 ES029213 and R01 ES025574 to JML and RJS.

## Acknowledgements

The authors thank Dr. Benjamin Laufer for WGBS computational biology expertise as well as collaborators in the MARBLES cohort (Dr. Irva Hertz Picciotto, Dr. Sally Ozonoff, Dr. Cheryl Walker) for human samples and data.

## Notes

https://github.com/cemordaunt/comethyl

https://www.ncbi.nlm.nih.gov/geo/query/acc.cgi?acc=GSE140730

### Competing Interest Statement

The authors have declared no competing interest.

