## Supplemental Figures for "Comethyl: A network-based methylome approach to investigate the multivariate nature of health and disease"

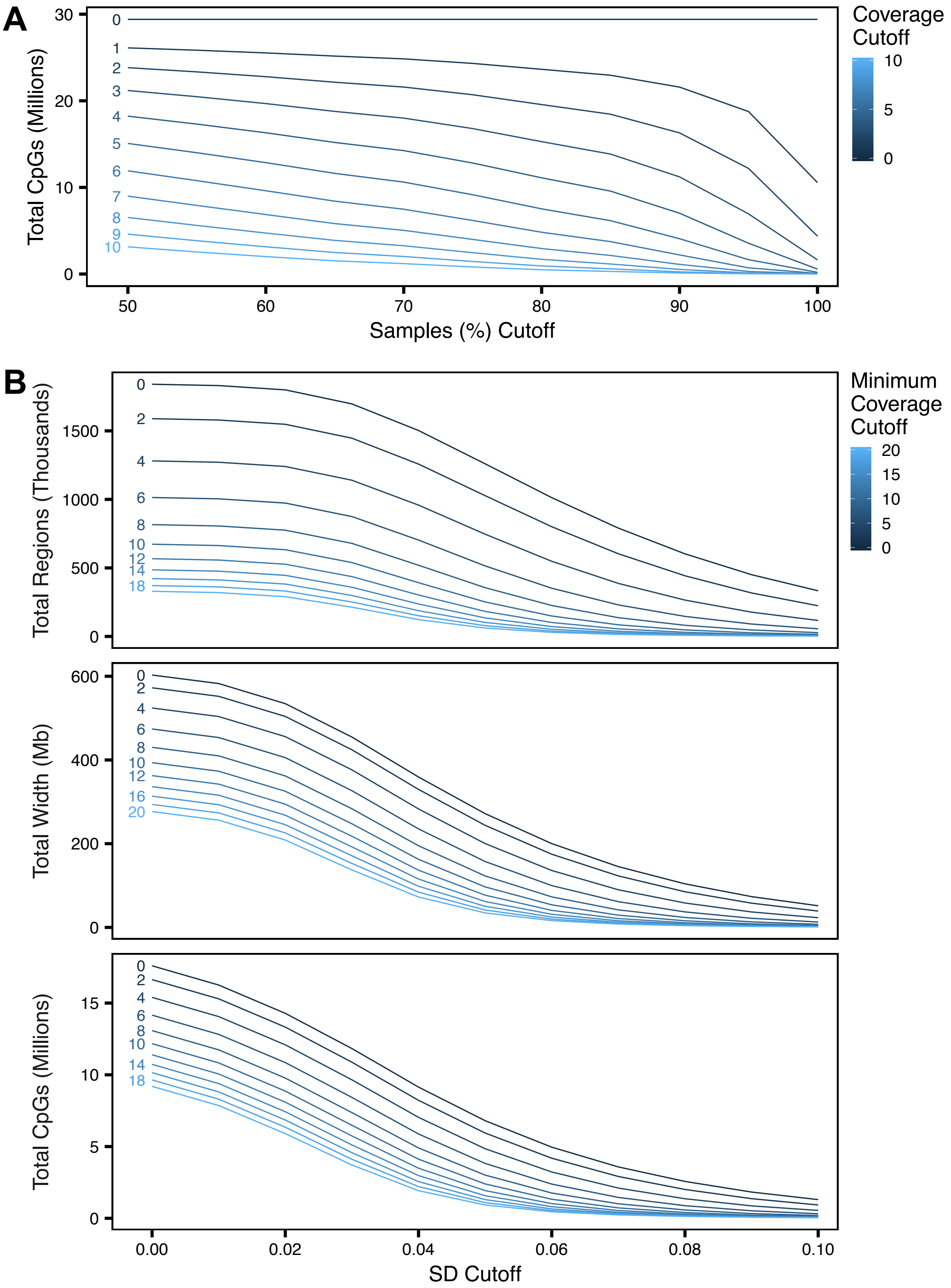

**Figure S1. CpG and region totals are altered by coverage and standard deviation filtering thresholds.** (A) Total CpGs passing various combinations of number of reads per sample and percent of samples with that number of reads. (B) Total regions, width, and CpGs included after filtering CpG cluster regions for different combinations of minimum reads in all samples and minimum methylation standard deviation.

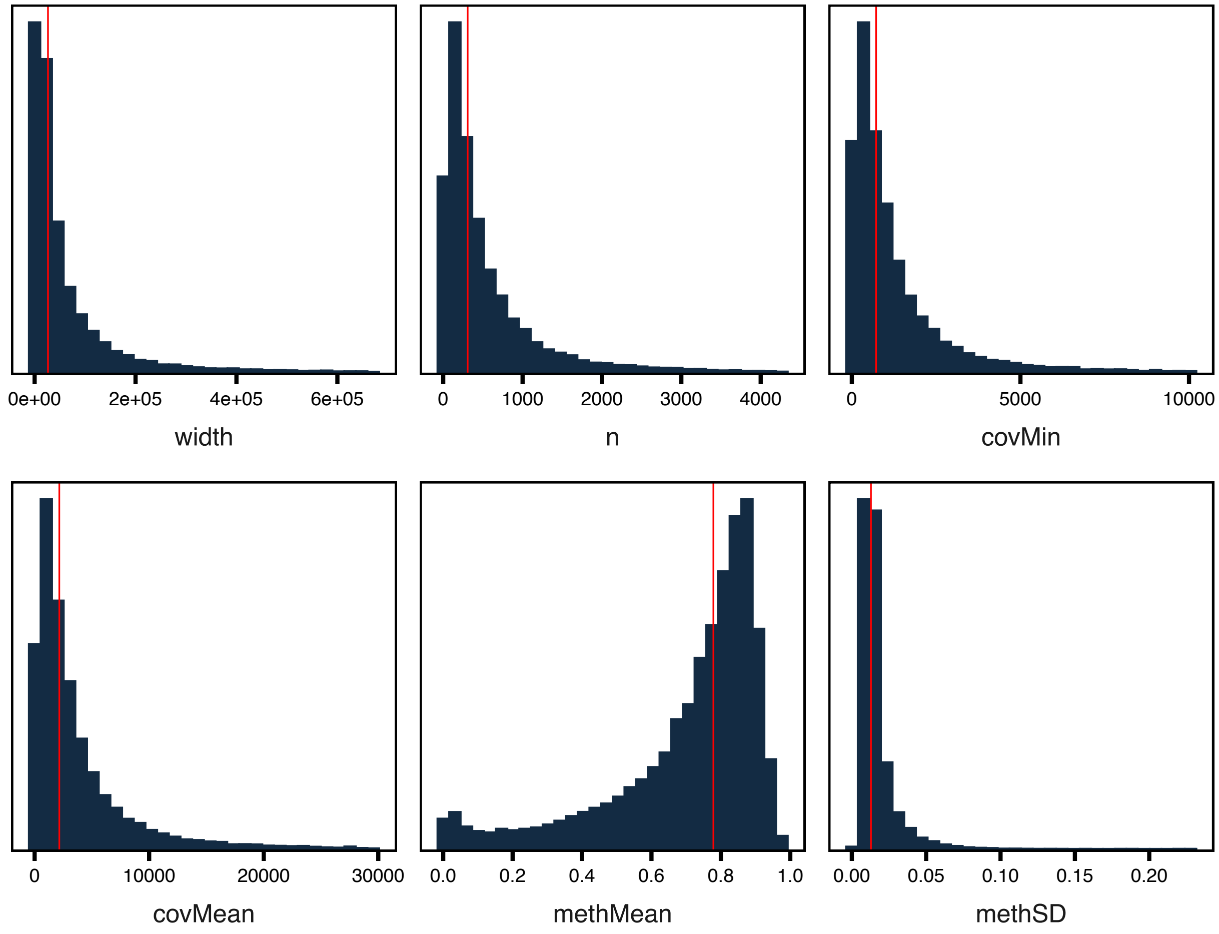


**Figure S2. After filtering of gene bodies, most regions have similar characteristics.** CpGs with sufficient coverage were grouped into regions defined by gene bodies and filtered for those with at least 3 CpGs and 10 reads in all samples. Plots show the distribution of multiple region characteristics with the red line indicating the median value.

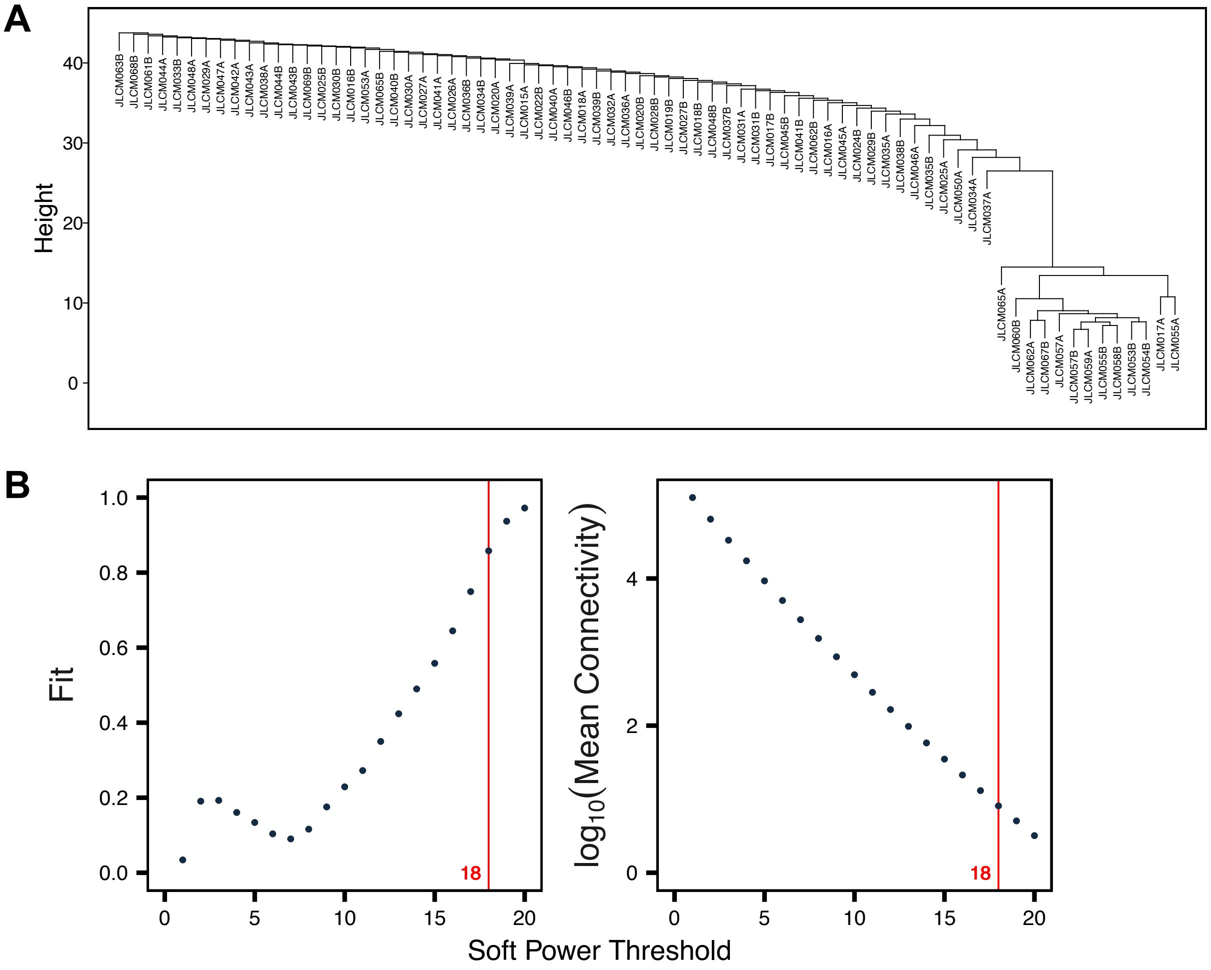


**Figure S3. CpG cluster-defined region methylation data from cord blood WGBS is appropriate for network construction.** Region methylation data was adjusted for the top 10 principal components and then examined for features relevant to network construction. (A) Samples were clustered by Euclidean distance to check for outliers. (B) Scale-free topology fit and mean connectivity is plotted at various soft power thresholds. The optimal soft power threshold of 18 for this dataset is shown in red.

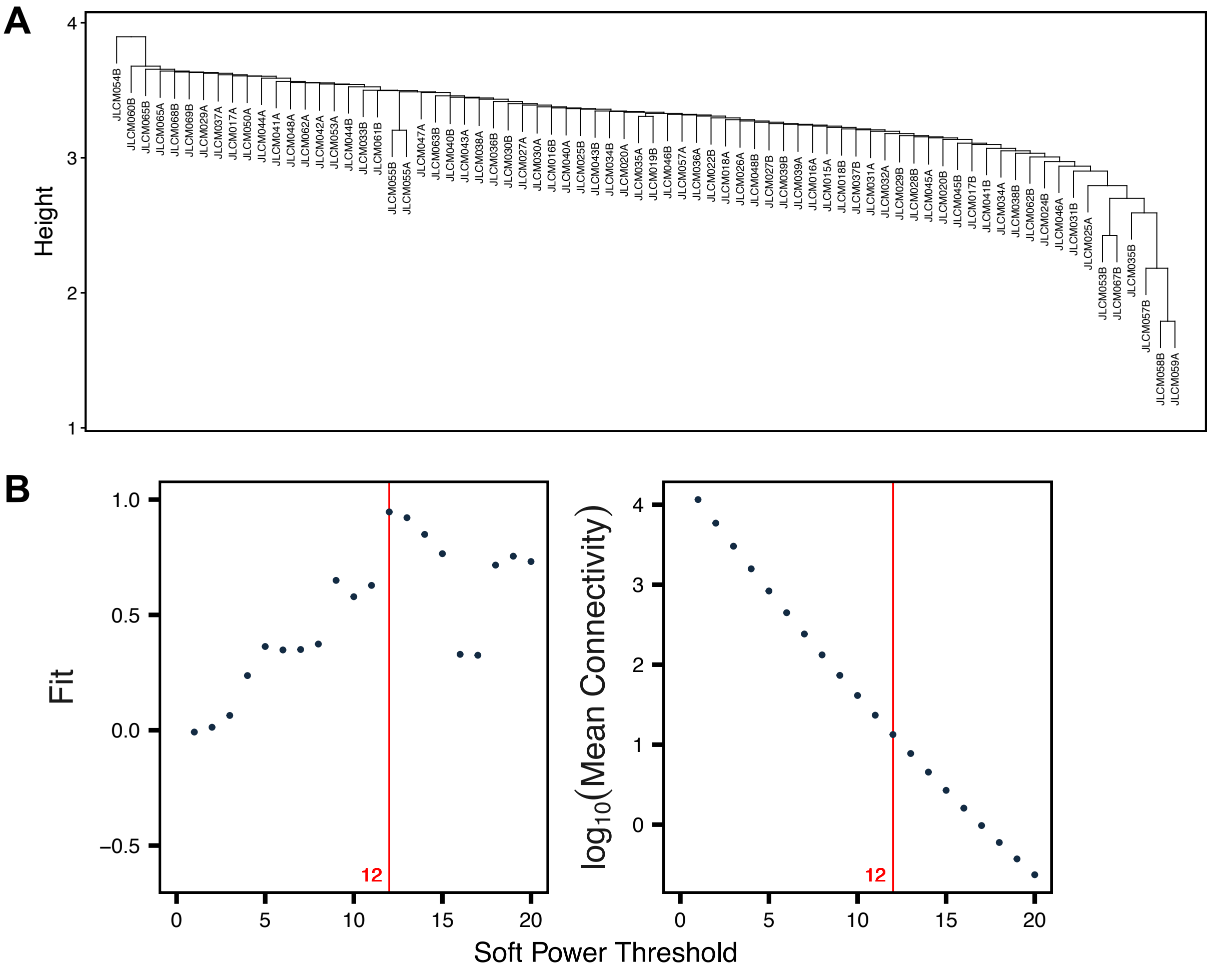


**Figure S4. Gene body-defined region methylation data from cord blood WGBS is appropriate for network construction.** Region methylation data was adjusted for the top 10 principal components and then examined for features relevant to network construction. (A) Samples were clustered by Euclidean distance to check for outliers. (B) Scale-free topology fit and mean connectivity is plotted at various soft power thresholds. The optimal soft power threshold of 12 for this dataset is shown in red.

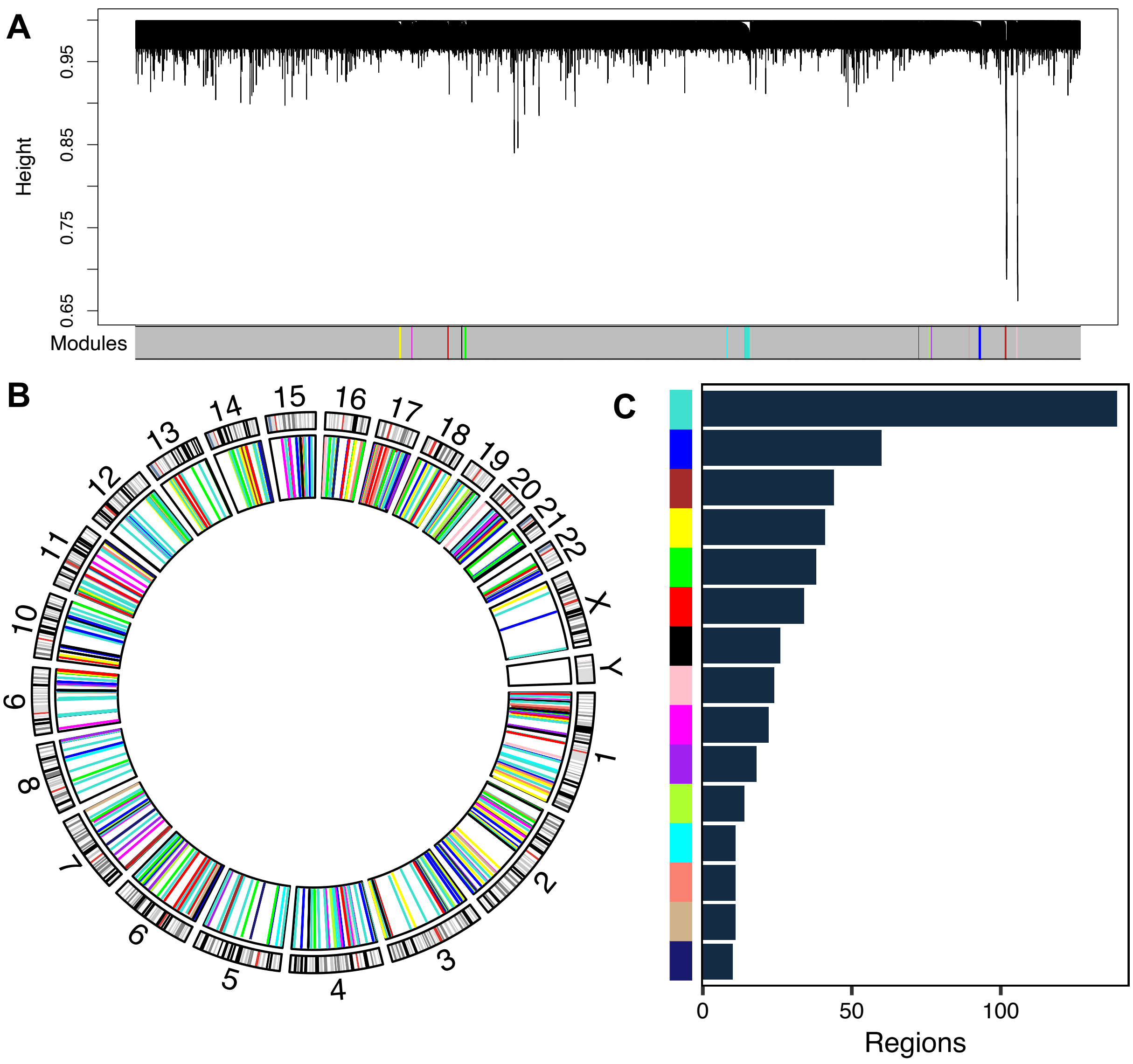


**Figure S5. Gene body comethylation modules identified in cord blood.** A comethylation network was constructed from adjusted region methylation data and assessed for modules of comethylated regions. (A) Region dendrograms for the only gene body region block. (B) Circos plot of regions colored by module in the inner ring and chromosome bands in the outer ring. (C) Number of regions assigned to each module.

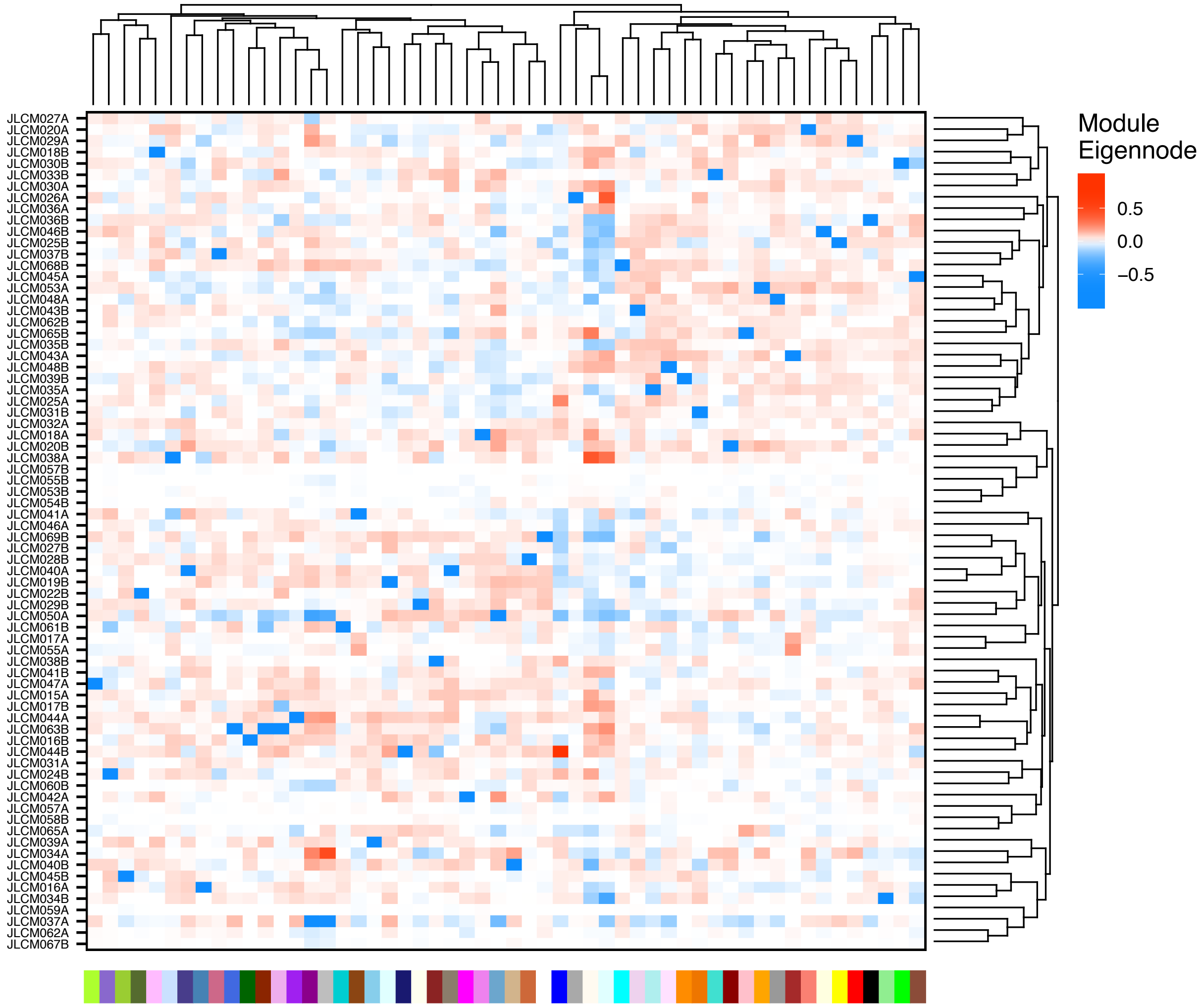


**Figure S6. CpG cluster module eigennode values for identified modules.** Samples and modules were clustered based on the biweight midcorrelation dissimilarity of the module eigennode values and the eigennode values were plotted in a heatmap.

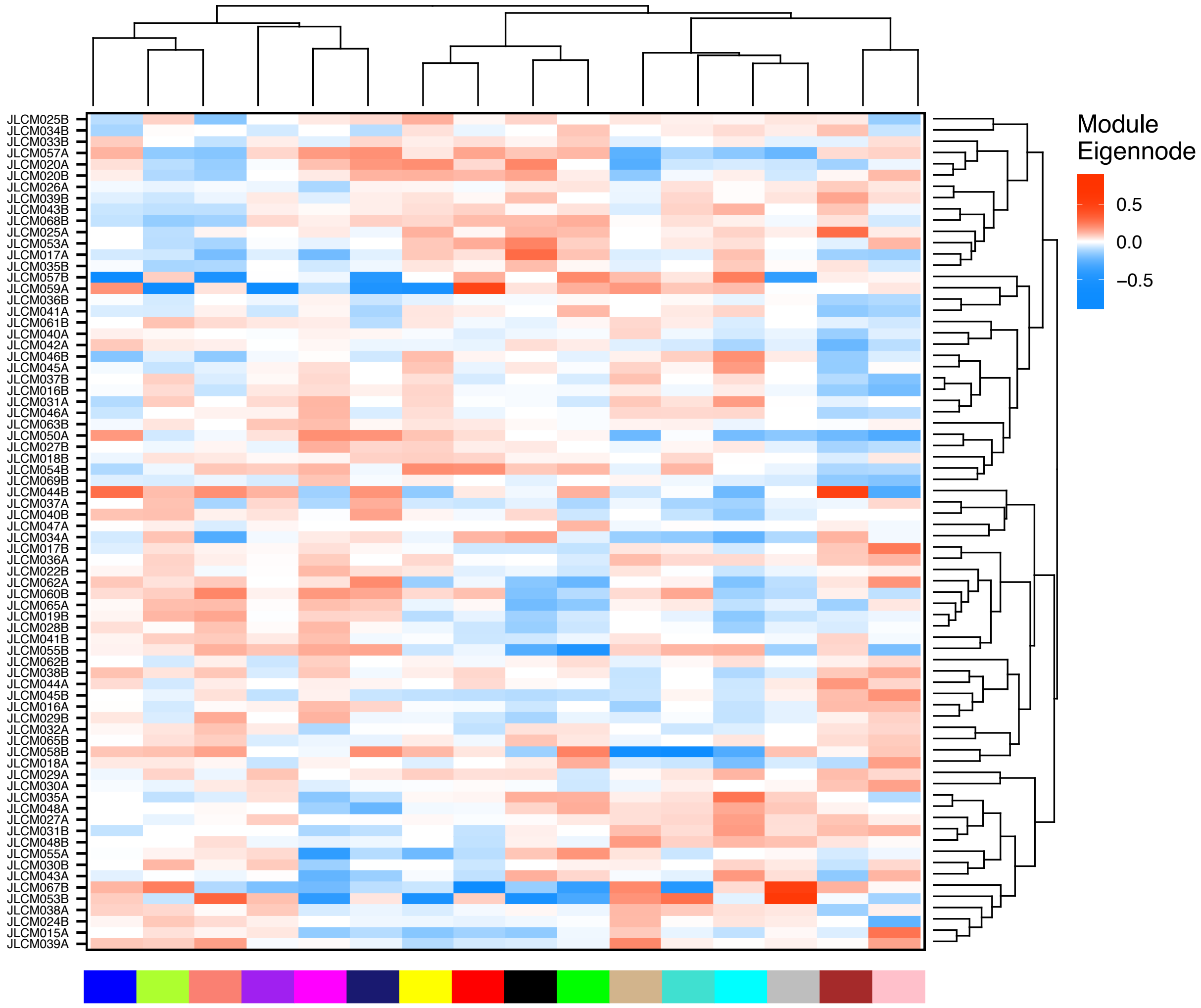


**Figure S7. Gene body module eigennode values for identified modules.** Samples and modules were clustered based on the biweight midcorrelation dissimilarity of the module eigennode values and the eigennode values were plotted in a heatmap.

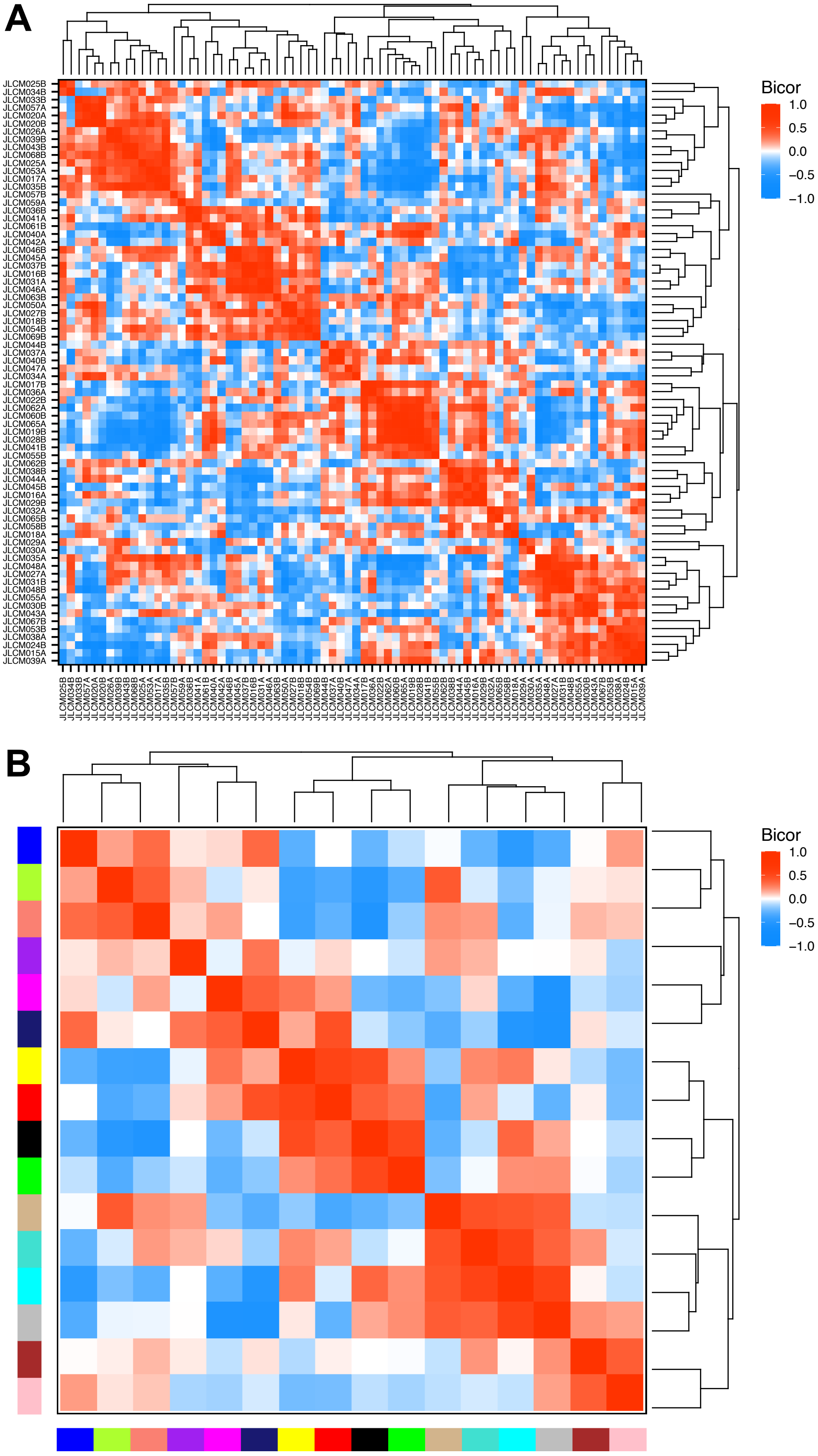
A
Bicor
1.0
0.5
0.0
−0.5
−1.0
B
Bicor
1.0
0.5
0.0
−0.5
−1.0

**Figure S8. Correlations of gene body module eigennode values reveal similarities between subsets of samples and modules.** (A) Eigennode values were clustered and compared across samples using biweight midcorrelation. (B) Same as (A), but compared across modules.

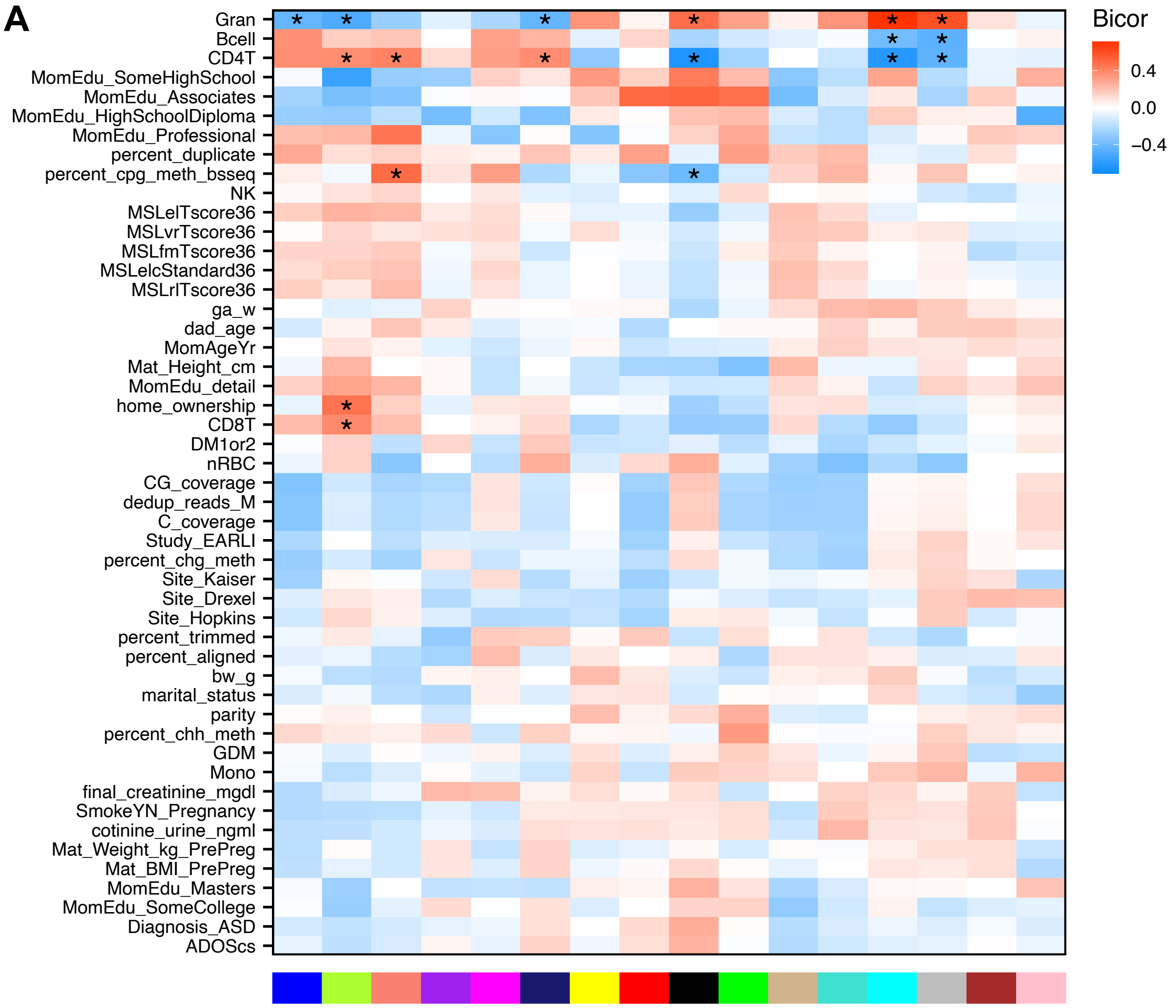


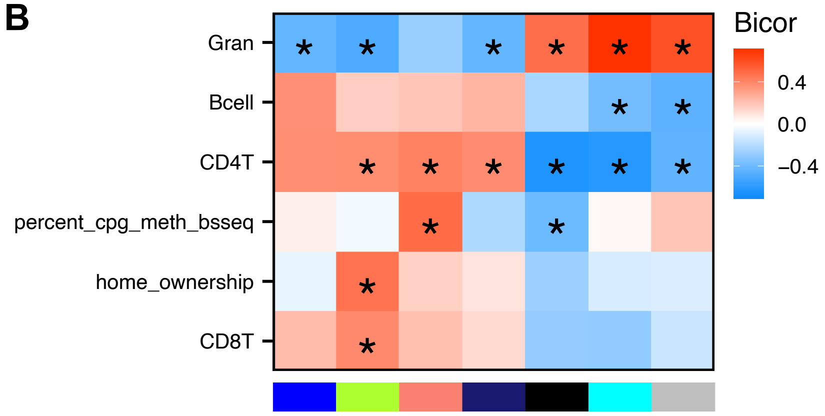


**Figure S9. Correlations of module eigennodes from gene body defined regions with sample traits.** (A) All tested correlations between module eigennode values and sample traits using biweight midcorrelation. (B) Same as (A) but significant modules and traits only (* FDR-adjusted $p < 0.05$).

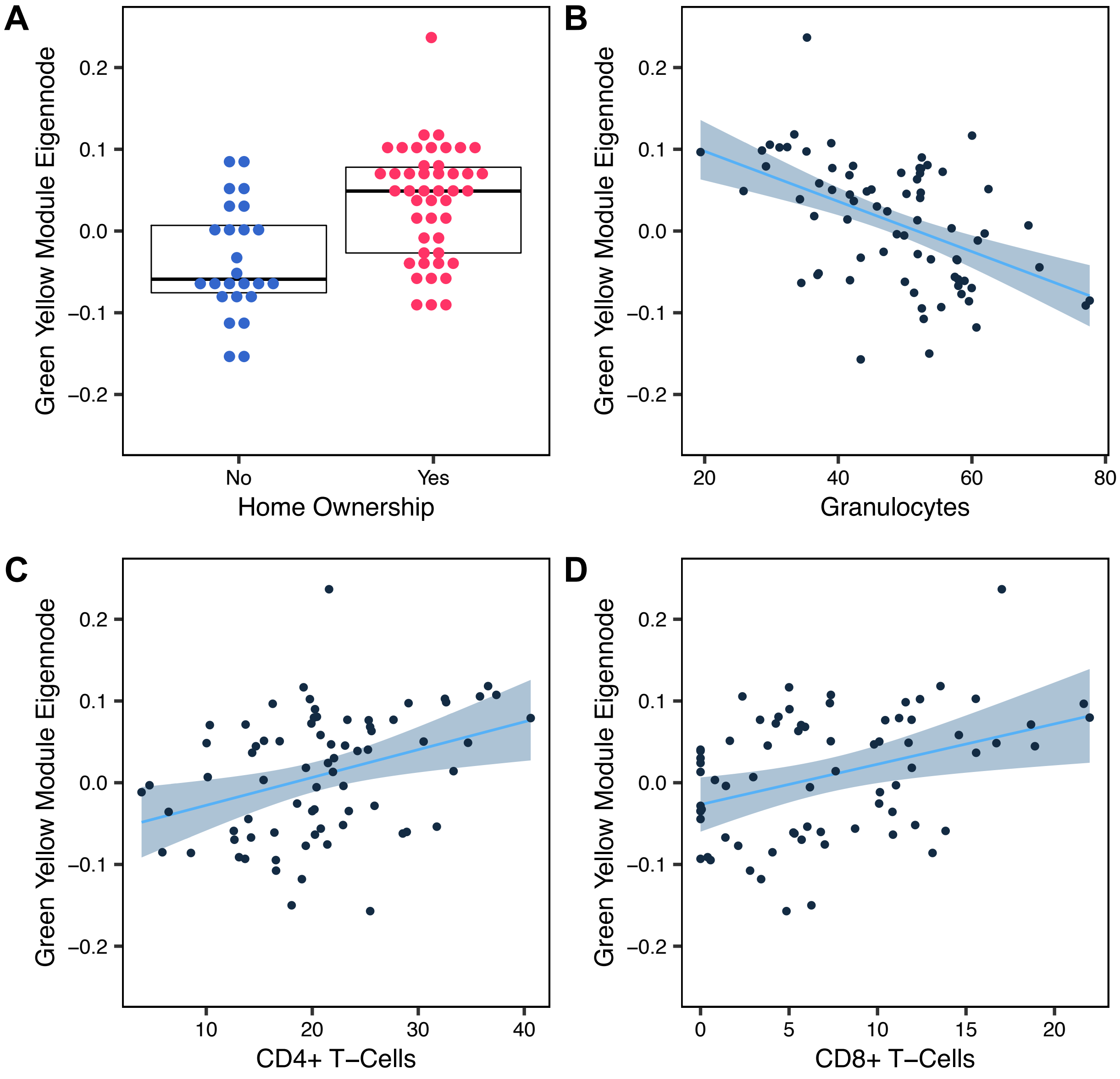


**Figure S10. The Green-Yellow Module from gene body regions is associated with home ownership and percentage of three cell types.** Eigennode values are plotted for each sample in relation to (A) home ownership, (B) granulocyte cell proportion, (C) CD4+ T-cell proportion, and (D) CD8+ T-cell proportion. Box in (A) indicates 1st quartile, median, and 3rd quartile. Lines in (B-D) are fit using robust regression and 95% confidence intervals are shaded.

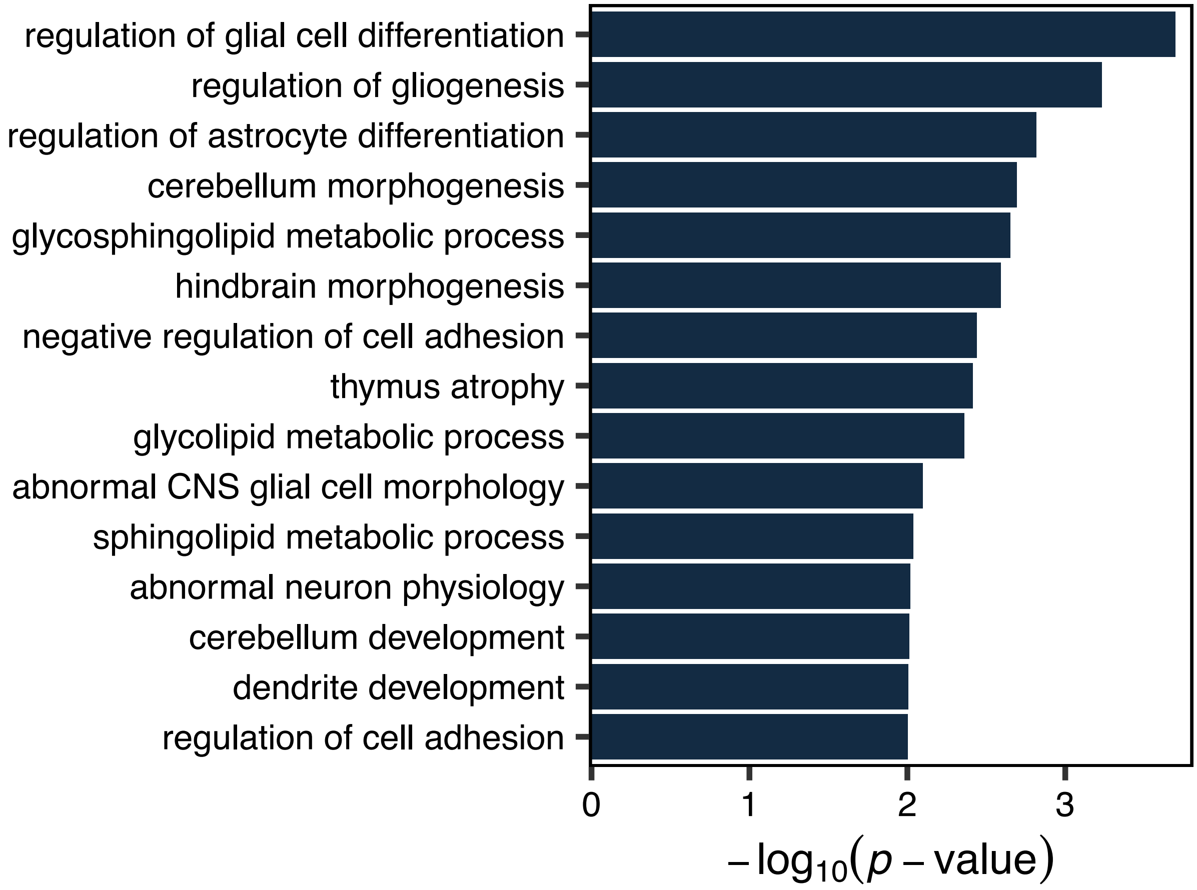


**Figure S11. Functional enrichment analysis of regions in the Bisque 4 Comethylation Module associated with later ASD diagnosis.** Enrichment analysis was performed using GREAT with Bisque 4 regions compared to all regions in the network for Gene Ontology, mouse phenotypes, and human phenotypes. Terms with an unadjusted *p*-value < 0.01 are shown.

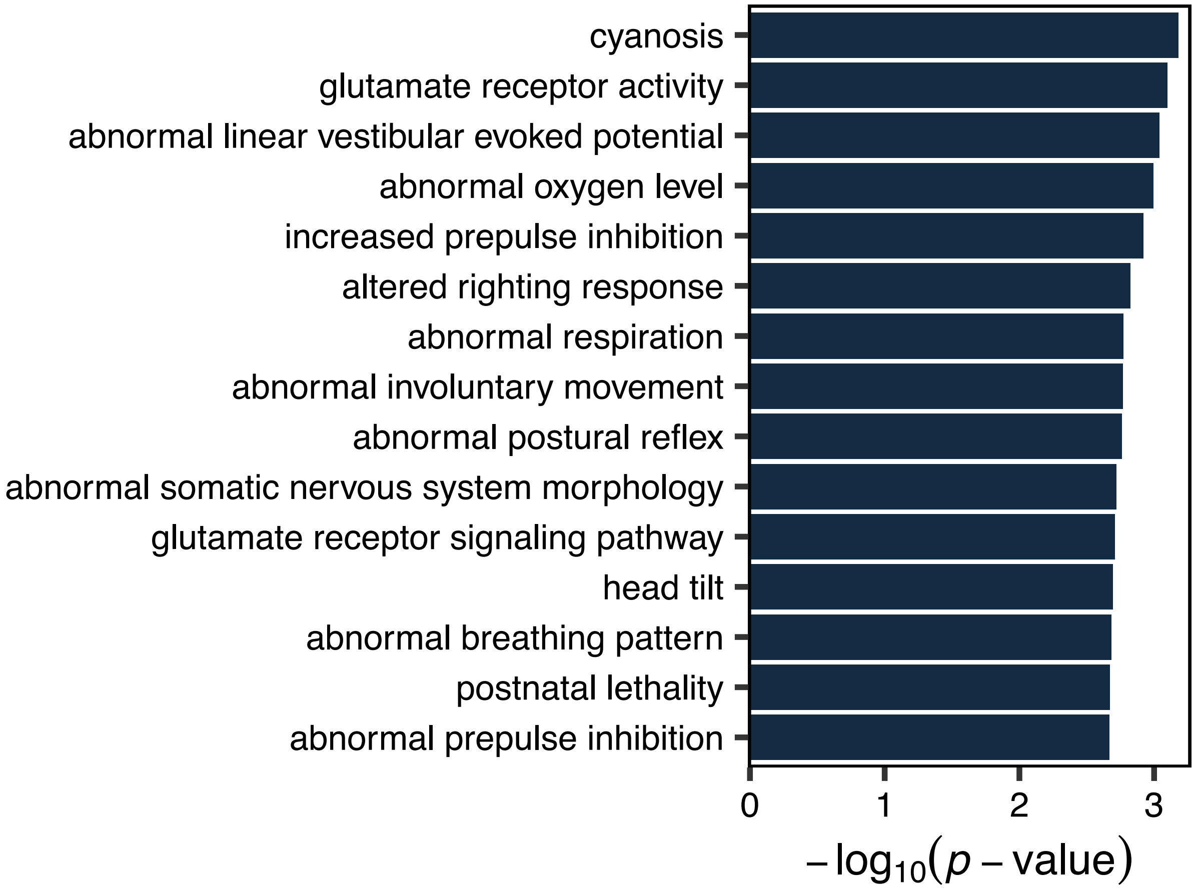


**Figure S12. Functional enrichment analysis of regions in the Green-Yellow Module from gene body analysis, associated with home ownership and 3 immune cell types.**

Enrichment analysis was performed using GREAT with Green-Yellow module regions compared to all regions in the network for Gene Ontology, mouse phenotypes, and human phenotypes. Terms with an unadjusted *p*-value < 0.01 are shown.
